# Inhibitor of apoptosis proteins determine glioblastoma stem-like cells fate depending on oxygen level

**DOI:** 10.1101/259283

**Authors:** Aurélie Soubéran, Jessica Cappaï, Mathieu Chocry, Christopher Nuccio, Julie Raujol, Carole Colin, Daniel Lafitte, Hervé Kovacic, Véronique Quillien, Nathalie Baeza-Kallee, Geneviève Rougon, Dominique Figarella-Branger, Aurélie Tchoghandjian

## Abstract

In glioblastomas, apoptosis inhibitor proteins (IAPs) are involved in apoptotic and non-apoptotic processes. Here we used GDC-0152, a small molecule IAP inhibitor, to explore how IAPs participate in glioblastoma stem-like cell maintenance and fate under both hypoxic and normoxic environments. In hypoxia, IAPs inhibition triggered stem-like cells apoptosis and decreased proliferation in four human glioblastoma cell lines, whereas in normoxia it induced a loss of stemness and differentiation. In addition, we characterized a 3D glioblastoma spheroid model. By using MALDI images we validated that GDC-0152 penetrates in the entire sphere. TOF-SIMS analyses revealed an oxygen gradient correlated with spatial cellular heterogeneity with proliferative and apoptotic cells located close to the hypoxic core and GFAP^+^ cells at the periphery. Notably, Serine-Threonine Kinases activation analysis revealed that oxygen level affects signaling pathways activated by GDC-0152. In hypoxia, IAPs inhibition activated ATR whereas in normoxia it activated NF-κB. Our data brings new mechanistic insights revealing the dual role of IAPs inhibitors like GDC-0152 that are relevant to their therapeutic application in tumors like glioblastomas.

## Introduction

Glioblastomas (GBs) are highly aggressive, infiltrative brain tumors. Numerous studies have shown that GBs are derived from cancer stem cells resistant to chemotherapy. Targeting GB stem cells which drive relapses and participate to treatment resistance represent a major challenge to improve patient’s overall survival. These GB stem cells are often located closed to hypoxic areas that maintain their self-renewal properties (Soeda *et al*, 2009), an environment that might also influence cells drug responses. They are characterized by cell surface antigens such as CD133 (Singh *et al*, 2004) or, as we have shown, gangliosides recognized by A2B5 antibody (Colin *et al*, 2006). A2B5^+^ GB cells harbor stem-like cell properties as they are able to self-renew to form spheres, proliferate and differentiate *in vitro* and initiate a tumor similar to the parental one *in vivo* (Tchoghandjian *et al*, 2010, 2012). In cancer cells, Inhibitor of Apoptosis Proteins (IAPs) are often overexpressed, they inhibit caspases activation and apoptosis, and therefore contribute to treatment resistance (Fulda & Vucic, 2012). In addition to controlling programmed cell death, IAPs regulate Mitogen-Activated Protein Kinase (MAPK) as well as both canonical and non-canonical Nuclear Factor-kB (NF-κB) pathways (Feltham *et al*, 2012; Varfolomeev *et al*, 2008; Bertrand *et al*, 2008; Vallabhapurapu *et al*, 2008; Zarnegar *et al*, 2008) leading to the transcription of target genes such as Tumor Necrosis Factor alpha (TNFα) (Varfolomeev *et al*, 2007; Vince *et al*, 2007; Tchoghandjian *et al*, 2013). IAPs expression level can be regulated by small molecules that mimic the N-terminal of Second Mitochondria-derived Activator of Caspase (Smac), an endogenous IAPs antagonist.

In a previous study we evaluated the prognostic value of the expression of cIAP1, cIAP2, XIAP and ML-IAP in human GBs and found that ML-IAP was correlated with the worst prognosis (Tchoghandjian *et al*, 2016). We used Smac mimetic GDC-0152 which antagonizes cIAP1, cIAP2, XIAP and also ML-IAP (Flygare *et al*, 2012) to test for its anti-tumoral activity in GBs. GDC-0152 treatment increased survival of mice xenografted with U87-MG GB cells while cultured GB stem-like cells were more resistant to GDC-0152-induced cell death (Tchoghandjian *et al*, 2016). We then explored whether IAPs affect GB stem-like cells differently depending on oxygen levels as hypoxia has been shown to modulate drug efficacy and IAPs expression (Dong *et al*, 2001; Yang *et al*, 2014; Hsieh *et al*, 2015). To this end, we compared IAPs inhibition by Smac mimetic GDC-0152 on GB stem-like cells differentiation, viability, proliferation and cell death under normoxic and hypoxic conditions.

## Results

### IAPs distribution in peri- and non-necrotic areas in glioblastomas

Intra-tumoral IAPs distribution within human GBs is not well documented. We sought to examine whether their distribution varied between hypoxic and non hypoxic regions of the tumors. We compared gene expression profiles of palisading cells of human GBs, representing the most hypoxic areas of tumors, to the other tumor cells (Dong *et al*, 2005). There was no significant difference of IAPs mRNA expression between palisading cells and the other tumor cells (Fig EV1A). These data showed that IAPs are homogeneously distributed within GBs independently of hypoxic areas.

### In vitro hypoxic conditions reflect human glioblastomas intra-tumoral microenvironment

In order to find a gene whose expression could discriminate hypoxic and non-hypoxic cells, we analyzed in the database cited before (Dong *et al*, 2005) the expression of HIF-1α target genes. We found that *adrenomedullin* (ADM), a gene known to be overexpressed in GBs, was significantly more expressed in palisading cells compared to the other tumor cells (Fig EV1B). Therefore we used ADM expression level as the read out of hypoxia. In our experiments to mimic GB microenvironment, GBM9 stem-like cells were cultivated either in normoxia (20% of O_2_) or hypoxia (2% of O_2_). After 8 days, ADM mRNA level was 15-fold increased in cells cultivated in hypoxia as compared to cells cultivated in normoxia, validating our hypoxic conditions (Fig EV1C). Then, we quantified mRNA expression levels of IAPs in GBM9 stem-like cells cultivated in these conditions. We found that IAPs mRNA expression levels were comparable between normoxia and hypoxia (Fig EV1D) confirming *in situ* results.

### IAPs inhibition by Smac mimetic GDC-0152 under normoxic and hypoxic conditions

In order to inhibit IAPs proteins expression, we treated four GB stem-like cells with Smac mimetic GDC-0152 both in normoxia and hypoxia. The expression was measured after 8 days of GDC-0152 treatment at two concentrations already shown to be not toxic in normoxia (0.01 nM and 1 nM) (Tchoghandjian *et al*, 2016) (Fig EV2). Results showed that 1 nM GDC-0152 decreased IAPs expression in GB stem-like cells, both in normoxia and in hypoxia. This concentration was chosen for the following experiments. Interestingly, in all GB stem-like cell lines tested, cells were more sensitive to the lower concentration of GDC-0152 under hypoxia as IAP proteins were more drastically decreased (*e.g*., in GBM9 cell line, cIAP1 was 1.4-fold decreased in hypoxia than in normoxia, XIAP 6-fold, cIAP2 1.15-fold and ML-IAP 1.2-fold upon 0.01 nM of GDC-0152). Based on these results, for the following experiments we aimed to target IAP proteins both in normoxia and hypoxia conditions. Therefore, we chose 1 nM of GDC-0152 as the effective concentration.

### IAPs inhibition impairs clonogenicity of glioblastoma stem-like cells only in normoxia

We previously described that Smac mimetics triggered GB stem-like cells differentiation into astrocytic GFAP^+^ cells (Tchoghandjian *et al*, 2014). Here we explored in four GB stem-like cell lines whether inhibiting IAPs with GDC-0152 differently alters the clonogenic potential and the expression of stem-like cell markers under normoxia or hypoxia. One nanomolar GDC-0152 pretreatment significantly reduced sphere formation by all cell lines: GBM6 (29%), GBM9 (38%), RNS175 (42%) and GBM40 (26%) but only in normoxia (Fig 1A and EV3A). Moreover IAPs inhibition decreased A2B5 expression in all GB stem-like cell lines (17% for GBM6; 15% for GBM9; 18% for RNS175; 5% for GBM40) and CD133 in GBM9 (46%) and GBM40 (55%) in normoxia only (Fig 1B-C,E and EV3B-C). In parallel, in normoxia, GDC-0152 treatment resulted in increased numbers of GFAP^+^ cells, which reflects differentiation, in three of the GB stem-like cell lines (171% for GBM6; 44% for GBM9; 33% for RNS175), whereas it did not alter GFAP expression in hypoxia (Fig 1D-E and EV3D). These results demonstrate that GDC-0152 modifies GB stem-like cells properties only in normoxia but not in hypoxia.

**Figure 1.**
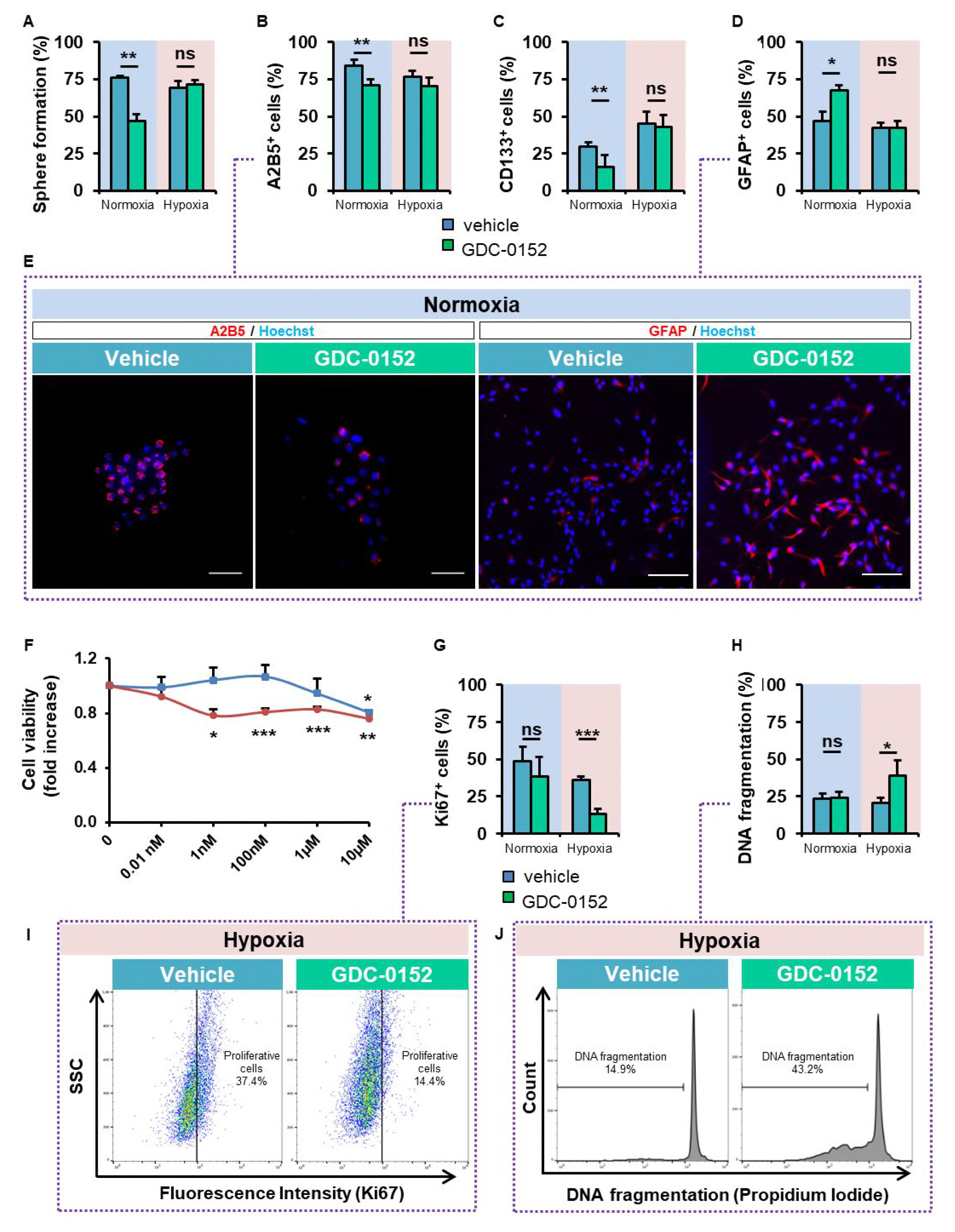
IAPs inhibition triggers loss of stemness in normoxia and decreases cell proliferation and increases cell death in hypoxia. (**A-E, G-J**) GBM9 cells grown in monolayers and treated with vehicle alone (DMSO) or 1 nM of GDC-0152 in normoxia or in hypoxia for 8 days. (**A**) Percentage of self-renewal calculated as the number of spheres formed divided by the number of cells seeded. Mean + SEM (*n*=3 independent experiments) is shown. **(B-D)** After treatment, cells were dissociated and stained either with A2B5 (*n*=8), anti-CD133 (*n*=8) or anti-GFAP (*n*=3) antibodies for flow cytometry analyses. Mean + SEM. **(E)** A2B5 (scale bars, 50 µm) and GFAP (scale bars, 100 µm) stainings of GBM9 stem-like cells counterstained with Hoechst. A representative staining of three experiments is shown. **(F)** GBM9 cells were treated in monolayer with increasing concentrations of GDC-0152 [0.01 nM; 1 nM; 100 nM; 1 µM; 10 µM] for 8 days in normoxia (blue) or hypoxia (red). Cell viability was expressed as fold increase of DMSO controls + SEM (*n*=4 in triplicate). **(G)** After treatment, cells were dissociated, fixed and stained with Ki67-antibody for flow cytometry analyses and the percentage of proliferation is shown for control and GDC-0152 treated cells for normoxia or hypoxia. Data are expressed as mean + SEM (*n*=4). **(H)** DNA fragmentation (SubG0/G1) determined by flow cytometry for control and GDC-0152-treated cells. Data are expressed as mean + SEM (*n*=5). **(I)** Representative dot plots of Ki67 flow cytometry of GBM9 cells from four independent experiments. **(J)** Representative histograms of DNA fragmentation of GBM9 from five independent experiments. Normoxia: 20% O_2_; hypoxia: 2% O_2_. \**P*<0.05; \*\**P*<0.01; \*\*\**P*<0.001; ns: not significant.

### IAPs inhibition decreases cell proliferation and increases apoptosis only in hypoxia

As we found that IAPs inhibition by GDC-0152 did not alter stem-like cells properties in hypoxia, we asked if it could affect GB stem-like cells’ viability. At 0.01 nM, GDC-0152 treatment in hypoxia significantly decreased cell viability of GBM6 (25%) and RNS175 (18%), and at 1 nM viability of all the four GB stem-like cell lines was affected (27.7% for GBM6; 21% for GBM9; 13.1% for RNS175; 20% for GBM40); neither concentration affected viability in normoxia (Fig 1F and EV3E). As cell viability reflects the balance between proliferation and apoptosis, we next quantified Ki67 expression, a marker of proliferation, and DNA fragmentation. IAPs inhibition decreased slightly cell proliferation in normoxia while in hypoxia, Ki67^+^ cells decreased drastically and significantly in the four GB stem-like cell lines (78% for GBM6; 64% for GBM9; 39% for RNS175 and 41% for GBM40; Fig 1G, I and EV3F). Morevover, IAPs inhibition triggered apoptosis only in hypoxia for all the GB stem-like cell lines (56% for GBM6; 86% for GBM9; 250% for RNS175 and 172% for GBM40; Fig 1H, J and EV3G).

Altogether these results demonstrate that IAPs inhibition alters cell viability by decreasing cell proliferation and increasing apoptosis in hypoxia but not in normoxia.

### Validation in a 3D glioblastoma model

In order to validate our results in an integrated GB 3D model we explored if GBM9 8-day-old spheres cultivated in normoxia reflect GB heterogeneity. To measure oxygen gradient, we performed TOF-SIMS analyses of secondary O^−^ ions. The data revealed the presence of a hypoxic core surrounded by a gradient of oxygen which increased from the core to the periphery of the sphere (Fig 2A). Analysis of cellular composition showed low rate PKH67^+^ proliferative cells localized in the core of the sphere (Fig EV4A).). Proliferative Ki67^+^ cells were located close to the hypoxic core whereas GFAP^+^ cells were found at the periphery of the spheres (Fig EV4B). By using MALDI images we demonstrated that GDC-0152 can penetrate into 8-day-old spheres in the entire sphere. The presence of GDC-0152 was confirmed by the colocalization of the [M+H]+ and the [M+Na]+ signal (Fig EV4C).

**Figure 2.**
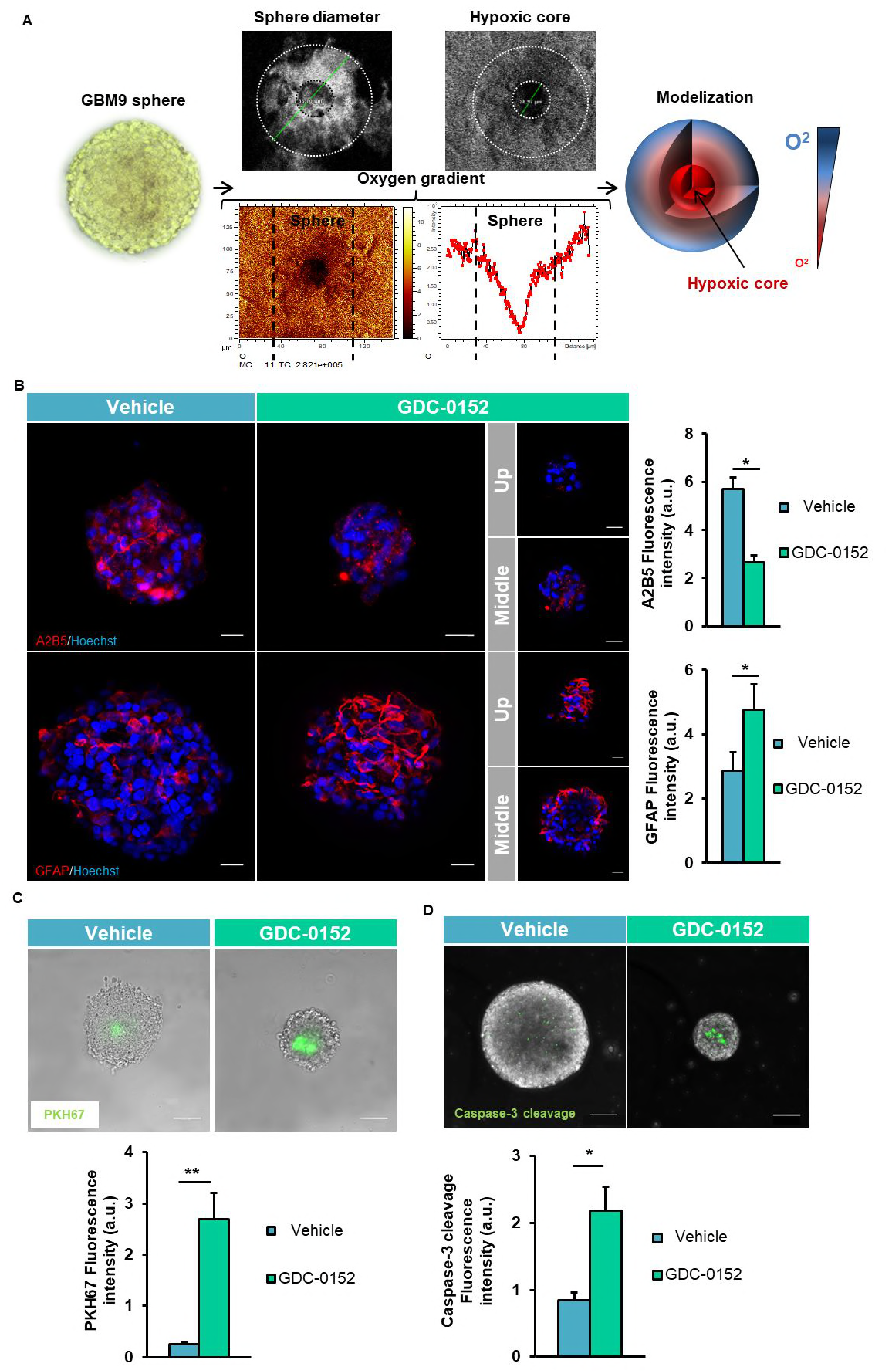
IAPs inhibition effects on a glioblastoma 3D model. (**A**) GBM9 cells were grown in suspension for 8 days in normoxia (20% O_2_). Spheres were analyzed by TOF-SIMS imaging. Sphere and hypoxic core diameters (left) and oxygen gradient (right) were measured. The amplitude of the color scale bar corresponds to the maximum number of count (MC). TC is the sum of counts recorded in all the pixels. Experiment was performed in triplicate. A modeling of spheres is presented on the right. (**B-D**) GBM9 cells were grown in suspension for 8 days and then treated with vehicle alone (DMSO) or GDC-0152 (1 µM) for 8 days. (**B**) GBM9 spheres were stained with A2B5 and anti-GFAP antibodies and counterstained with Hoechst. Representative pictures of 15 z stacks projection (*n*=3 independent experiments) are shown. (**C**) GBM9 spheres were grown with PKH67 then green fluorescence was quantified after 8 days of treatment. (**D**) To monitor apoptosis in viable spheres, spheres were incubated with fluorescent caspase-3. (**B**-**D**) Quantification of fluorescence intensity was performed by ImageJ after 8 days of treatment (n=5 spheres) and divided by the size of the sphere (*n*=3 independent experiments). Scale bar, 20 µm.

This 3D GB model which mimics GBs heterogeneity, cellular organization and composition, was therefore suitable to analyze IAPs inhibition in relation with oxygen levels.

GDC-0152 treatment of spheres resulted in a decreased of A2B5^+^ cells of approximately 52% and an increase of GFAP^+^ cells of approximately 56% (Fig 2B). Moreover, PKH67 staining and caspase-3 cleavage quantified in the core of the GDC-0152-treated spheres, revealed a decrease in proliferation and an induction of apoptosis, respectively, in the most hypoxic areas of the spheres (Fig 2C-D). Taken together, these results validate in a 3D model that IAPs inhibition triggers differentiation in the most oxygenated areas, and decreases proliferation whilst promoting apoptosis in the most hypoxic zones of the spheres.

### Differential IAPs inhibition effects are supported by different cell signaling pathways

To dissect the specific pathways triggered by IAPs inhibition depending on oxygen level, we performed high throughput screening of Serine-Threonine Kinases (STK) activity in GBM9 stem-like cells either treated with DMSO or GDC-0152 at 1 nM for 2 h in normoxia and in hypoxia. Only STK more activated (negative values) or less activated (positive values) by GDC-0152 were considered and kinases involved in hypoxia response alone were removed from the analysis (Fig EV5). MAPK, JNK, PI3K/Akt and cell cycle pathways were found as the most commonly represented and activated both in normoxia and in hypoxia upon GDC-0152 treatment. In comparison, NF-κB and ATR pathways were differentially activated upon GDC-0152 treatment as represented (Fig 3A and Fig EV5). In normoxia GDC-0152 increased the activation of IKKb and RSK1,4 which enhanced NF-κB signaling, as usually found with Smac mimetic treatment, and it increased the inhibition of Chk1 which is the readout of ATR activity. In contrast, GDC-0152 increased ATR activity only in hypoxia. To validate these results, we analyzed phospho-IκBα, phospho-ERK1/2 and phospho-Chk1 proteins expression in GBM9 stem-like cells after 8 days of GDC-0152 treatment or DMSO. IAPs inhibition triggered phosphorylation of IκBα only in normoxia, phosphorylation of Chk1 only in hypoxia and phosphorylation of ERK1/2 in both conditions (Fig 3B-C).

**Figure 3.**
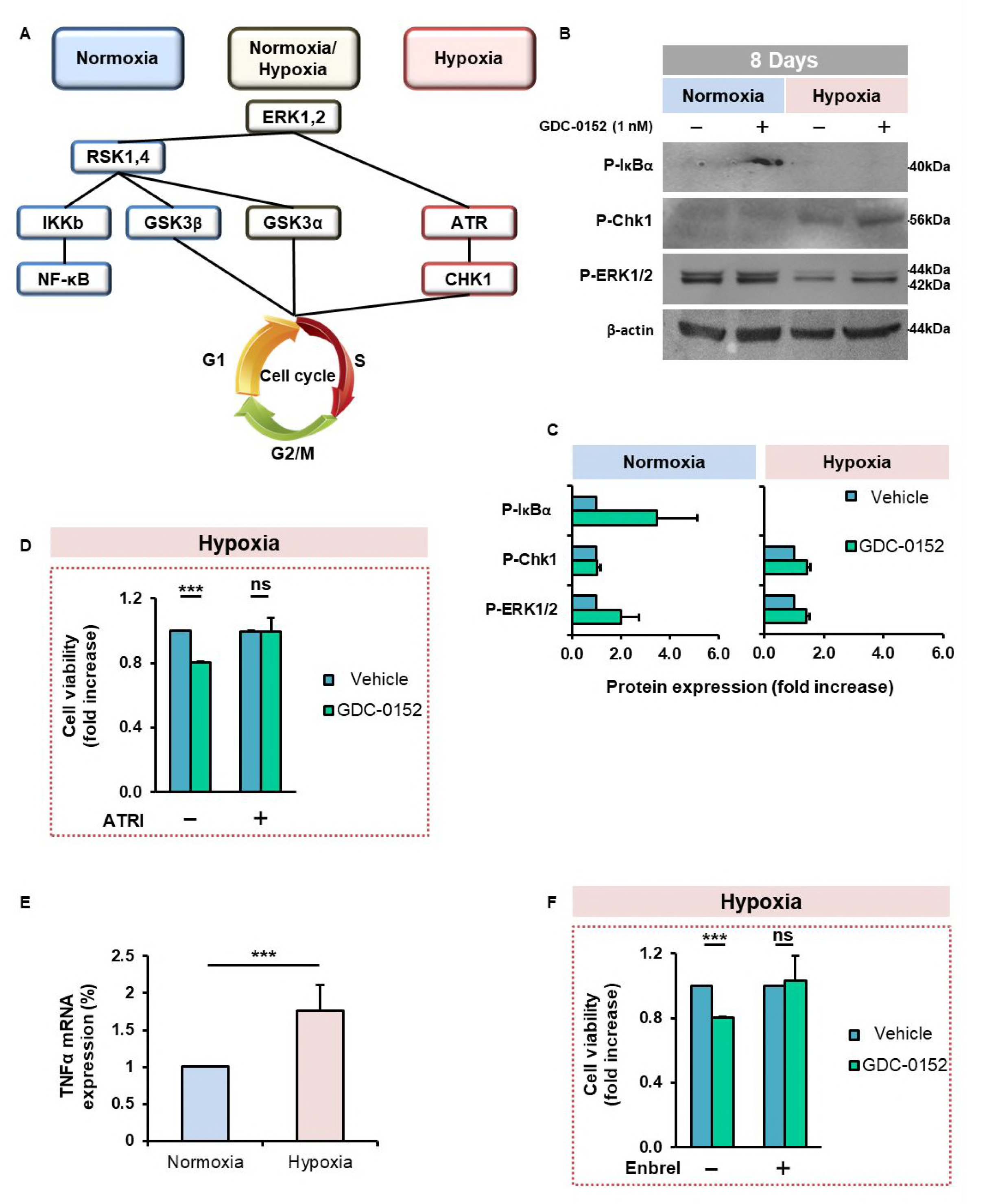
Signaling pathways involved upon IAPs inhibition depending on oxygen level. (**A**) GBM9 stem-like cells were cultivated in monolayer in normoxia or hypoxia and treated with vehicle alone (DMSO) or 1 nM of GDC-0152 for 2 h. Serine/threonine kinases activity was analyzed by PamGen kinome assay. The scheme represents the signaling pathways triggered by IAPs inhibition depending on microenvironment. (**B**) Validation of kinome results on GBM9 stem-like cells at 8 days of vehicle or 1 nM of GDC-0152 treatment in normoxia or in hypoxia. Phospho(P)-IκBα for NF-κB pathway, phospho(P)-ERK1/2 for MAPK pathway and phospho(P)-Chk1 were analyzed by western blotting. (**C**) Quantification of western blot analyses *(n =* 2). (**D**) Cell viability of GBM9 stem like-cells cultivated in hypoxia, treated with 1 nM of GDC-0152 alone or co-treated with GDC-0152 and 10 nM of ATR inhibitor (ATRI) was measured by MTT assay and expressed as fold increase of ATRI controls + SEM (*n*=3 independent experiments). (**E**) TNFα mRNA level was analyzed by Q-RT-PCR and fold increase TNFα mRNA level is shown + SEM (*n*=3 independent experiments). (**F**) Cell viability of GBM9 cells cultivated in hypoxia, treated with 1 nM of GDC-0152 alone or co-treated with GDC-0152 and 25 µg/ml of Enbrel (TNFα blocking antibody) was measured by MTT assay and expressed as fold increase of Enbrel controls + SEM *(n=4* independent experiments). Each experiment was performed in triplicate. \*\*\**P*<0.001. ns: not significant.

To test whether in hypoxia IAPs inhibition decreased cell viability through an ATR-dependent manner, we used a specific inhibitor of ATR activity (ATRI) in co-treatment with GDC-0152. Results showed that in combination with ATRI, effect of IAPs inhibition on cell viability was totally reversed (Fig 3D).

Moreover, we quantified TNFα mRNA expression and found it significantly more expressed in hypoxia than in normoxia (Fig 3E). Then, we used TNFα blocking antibody Enbrel in combination with GDC-0152. TNFα inhibition completely blocked the effect of IAPs inhibition on cell viability in hypoxia (Fig 3F).

Taken together, these results demonstrate that IAPs inhibition triggers different cell signaling pathways depending on oxygen level. In normoxia, IAPs inhibition activates NF-κB pathway whereas under hypoxia it activates ATR and TNFα signaling is involved.

## Discussion

This study shows that inhibition of IAPs expression by Smac mimetic GDC-0152 determines GB stem-like cells fate. In an environment rich in oxygen, GB stem-like cells lose their stem cell properties. In contrast, in an environment deprived of oxygen, IAPs inhibition does not affect stemness but rather cell viability by decreasing proliferation and increasing apoptosis.

We showed that these dual effects due to IAPs inhibition are driven by distinct signaling pathways. In normoxia, IAPs inhibition triggers the loss of stem cell properties and NF-κB is activated confirming our previous study (Tchoghandjian *et al*, 2014). In hypoxia, IAPs inhibition activates ATR as Smac mimetic treatment triggers phosphorylation of its principal target Chk1 and pharmacological inhibition of ATR blocks the effect of IAPs inhibition on cell viability. ATR, one of the main proteins involved in the DNA Damage Response, is essential for the maintenance of genomic integrity during a broad spectrum of DNA damages; it controls cell cycle and cell survival even in the absence of DNA damage (e.g. replicative stress; (Murga *et al*, 2009; Shiotani *et al*, 2013)). In this study, ATR was activated after only 2 h of IAPs inhibition in hypoxia suggesting an activation independent of any DNA damage.

Toledo *et al.* proposed a tumor-suppressive potential of ATR activation in a DNA-damage-independent manner by promoting cell cycle arrest (Toledo *et al*, 2008). As ATR activation has also been described to trigger apoptosis, in our conditions ATR could be involved both in proliferation arrest and apoptosis induction (Wei *et al*, 2016; García *et al*, 2015). Moreover, we showed that TNFα expression was increased in hypoxia and contributed also to IAPs inhibition-decreased cell viability in hypoxia as the use of Enbrel impaired Smac mimetic effect on cell viability. TNFα is well known to potentiate Smac mimetics efficiency (Lalaoui *et al*, 2016) and probably synergizes with GDC-0152 in hypoxia to trigger cell death. However, how and whether TNFα and ATR pathways are connected or act in parallel remains to be determined.

We found that Smac mimetic GDC-0152 decreases more drastically IAP proteins expression in hypoxia and at lower concentration than in normoxia. Our findings are consistent with those of Lu *et al*. on the Smac mimetic AT-406 in cervical cancer. They showed that upon AT-406 treatment, cells were more sensitive to radiation in hypoxia than in normoxia and that cIAP1 and XIAP degradation was increased in hypoxia (Lu *et al*, 2014). These results support the implication of a direct endogen IAP protein antagonist which could be down-regulated in hypoxia similarly to Smac or deubiquitinases (Guo *et al*, 2014; Lee & Song, 2015; Engel *et al*, 2016).

The cyto-architecture of GBs is complex and covered by a gradient of oxygen. The use of GB organoïds described by others (Hubert *et al*, 2016) was not suitable for our purpose. Therefore, we characterized and used an integrated 3D GB model to mimic GBs cellular complexity and physiology allowing to quantify, compare and correlate IAPs inhibition effects with oxygen level. The sphere model that we set up is instrumental to analyze drug responses in terms of phenotypic remodeling, apoptosis and proliferation in parallel with oxygen level variations.

Eradication of tumor stem cells is a major current challenge. Here we describe that IAPs inhibition *via* the use of a Smac mimetic determines stem cells fate by targeting them whatever the oxygen level. Hypoxia is a major cause of treatment resistance in solid cancer. In a hypoxic environment, cells gain stemness and become more resistant to conventional therapy. Therefore, the identification of this dual role of IAP proteins, and thereby of Smac mimetics, is of high interest for clinical purpose, in particular to target stem cells located in hypoxic areas of solid tumors.

## Materials and Methods

### Microarray data source

Expression profile by microarrays was obtained from National Center for Biotechnology Information Gene Expression Omnibus (NCBI GEO, http://www.ncbi.nlm.nih.gov/geo/) (Edgar *et al*, 2002; Barrett *et al*, 2013). We downloaded and re-analyzed the raw data of a previous Affymetrix Human Genome U133 Plus 2.0 Array (GDS1380) study that investigated molecules differentially expressed in laser capture microdissected peri-necrotic palisades *versus* other GB tumor area (Dong *et al*, 2005). We focused on the differential transcriptomic expression profiles of cIAP1 (202076_at), cIAP2 (210538_s_at), XIAP (206536_s_at) and ML-IAP (220451_s_at) in these two groups of samples.

### Cell lines and reagents

Four primary GB stem-like cell lines were used. GBM6, GBM9 and GBM40 were isolated from different human GBs and derived from A2B5^+^ cells (Tchoghandjian *et al*, 2012) and RNS175 was isolated from human GB without any selection. All these cell lines are *IDH*^wt^. These cells were grown as floating spheres in serum-free medium supplemented with EGF, bFGF and B27 (Tchoghandjian *et al*, 2010). Smac mimetic GDC-0152 was purchased from Selleckchem (Houston, Tex., USA) and ATR inhibitor from Merck KGaA (ATRI III; ETP-46464, Darmstadt, Germany). TNFα blocking antibody Enbrel (Pfizer, N.Y., USA) was kindly provided by S. Guis. For GDC-0152 treatment experiments, cells were grown either as spheres in suspension or as monolayers on 10 μg/ml poly-DL-ornithine (Sigma-Aldrich) coated dishes. Spheres were treated after 8 days of culture for 8 days whereas cells cultivated in monolayer on 10 μg/ml poly-DL-ornithin (Sigma-Aldrich) were treated after 24 h for 8 days. All the cell lines were grown at 37°C in a humidified atmosphere of 5% CO_2_ and 95% air including 20% O_2_ for normoxic condition or in a humidified atmosphere of 5% CO_2_ and 2% O_2_ supplemented with nitrogen (trigaz incubator O_2_-N_2_-CO_2_, Sanyo, Osaka, Japan) for hypoxic condition.

### Reverse-Transcription, Real-Time Quantitative PCR Analysis

Total RNA was extracted using E.Z.N.A blood RNA kit (Omega, Norcross, Ga., USA) according to the manufacturer’s instructions. Reverse Transcription was performed with SuperScript RT II (Invitrogen, Carlsbad, Calif., USA) according to manufacturer instructions, at 42°C for 2 h. Ribosomal 18S, glyceraldehyde-3-phosphate-dehydrogenase (GAPDH) and β-actin were used as reference genes. cIAP1, cIAP2, XIAP, ML-IAP, ADM and TNFα transcripts were analyzed by quantitative RT-PCR using a LightCycler® 480 and LightCycler 480 SYBRGreen I Master (Roche Applied Science, Meylan, France). The relative expression ratio of the target mRNA and reference RNA (18S, GAPDH, β-actin) was calculated using Q-PCR efficiencies and the crossing point Cp deviation of a stem-like cell lines *versus* normoxic DMSO control. All determinations were performed in triplicate. Results are expressed as median. Forward and reverse primers for each gene are listed below *18S:* 5’-CTACCACATCCAAGGAAGGCA-3’, 5’-TTTTTCGTCAACTACCTCCCCG-3’; *GAPDH:* 5’-CAAATTCCATGGCACCGTC-3’, 5’-CCCACTTGATTTTGGAGGGA-3’; β*-actin:* 5’-CCACACTGTGCCCATCTACG-3’, 5’- AGGATCTTCAATGAGGTAGTCAGTCAG-3’; *cIAP1:* 5’- CTGGCCATCTAGTGTTCCAG-3’, 5’-TCTACCCATGGATCATCTCC-3’; *cIAP2:* 5’-CTGCTATCCACATCAGACAG-3’, 5’-CCAGGCTTCTACTAAAGCCC-3’; *XIAP:* 5’-GGGGTTCAGTTTCAAGGAC-3’, 5’-GCGCCTTAGCTGCTCTTCAG-3’; *ML-IAP*: 5’-CCTGACAGAGGAGGAAGAGG-3’, 5’-ACCTCACCTTGTCCTGATGG-3’; *ADM*: 5′-TGCCCAGACCCTTATTCGG-3’, 5′-AGTTGTTCATGCTCTGGCGG-3′ and *TNFα*: 5’-ACAACCCTCAGACGCCACAT-3’, 5’-TCCTTTCCAGGGGAGAGAGG-3’.

### Protein extraction and western blotting

Proteins were extracted with RIPA lysis buffer supplemented with protease (Roche Applied Science) and phosphatase (Santa Cruz, for phospho-proteins) inhibitors for 30 min on ice followed by 10 min centrifugation at 12000 rpm. Fifty micrograms of proteins per lane were separated by 12% SDS-PAGE and transferred onto nitrocellulose membrane. After 1 h of blocking in 5% skimmed milk, membranes were incubated with the following antibodies: anti-cIAP1 (0.2 µg/ml, R&D Systems, Minneapolis, Minn., USA), anti-cIAP2 (0.2 µg/ml, Merck KGaA), anti-XIAP (0.25 µg/ml, clone 28, BD Biosciences, Franklin Lakes, N.J., USA), anti-ML-IAP (0.5 µg/ml, clone 88C570, Novus, St. Louis, Mo., USA), anti-β-actin (1/5000, clone AC-15, Merck KGaA), anti-Phospho-Chk1 (Ser345) (1:1000, 2348 clone 133D3), anti-Phospho-ERK1/2 (1:2000) and anti-Phospho-IκBα (1:1000) all from Cell signaling technology (Danvers, Mass., USA) in TBS supplemented with 5% BSA and 0.1% tween 20 (Sigma-Aldrich) overnight at 4°C under shaking. The following horseradish peroxidase-conjugated donkey anti-mouse IgG, goat anti-rabbit IgG (Santa Cruz Biotechnology, Dallas, Tex., USA) and rabbit anti-goat IgG (Dako France, Ulis, France) antibodies were used for the detection of IAP proteins. The following fluorescent secondary antibodies were used: donkey anti-rabbit-800CW or donkey anti-mouse-680LT (LI-COR, Lincoln, Neb., USA) and detected with Odyssey® CLx Imaging System from LI-COR for the phospho-proteins.

### Flow cytometry

Anti-CD133-PE (CD133/2) and A2B5-APC antibodies were purchased by Miltenyi Biotec (Bergisch Gladbach, Germany); staining assays were performed as previously described (Tchoghandjian *et al*, 2010). For GFAP and Ki67 stainings, cells were harvested, fixed at room temperature in 2% paraformaldehyde for 10 min. Then cells were permeabilized with denaturating buffer (HCl 37%, Triton X100, PBS 10X) for 20 min at 37°C and neutralized with Sodium tetraborate for 10 min at room temperature. Cells were incubated with anti-GFAP-PE (BD biosciences) and anti-Ki67-FITC (clone MIB1, Dako) for 30 min at room temperature. Cells were processed on a FACS Calibur (Becton Dickinson, Heidelberg, Germany). Data were analyzed using FlowJo software (Tree Star, Inc., Ashland, Or., USA).

### DNA fragmentation

Fluorescence-activated cell sorting (FACS Calibur) analysis of DNA fragmentation of propidium iodide-stained nuclei was performed as described (Tchoghandjian *et al*, 2016).

### Immunofluorescence

Before GFAP and Ki67 staining, cells or spheres were fixed with 4% paraformaldehyde, denaturated (HCl 37%, Triton X100, PBS 10X) for 20 min at 37°C and neutralized with Sodium tetraborate for 10 min at room temperature. Primary antibodies A2B5 (mouse IgM ascite, 1:1000, kindly provided by G. Rougon), anti-GFAP (rabbit IgG, 10 µg/mL, Dako) and anti-Ki67 (mouse IgG, clone Mib1, 5 µg/ml, Dako) were incubated for 1 h at room temperature. For A2B5, cells were fixed after staining. Double staining assays, GFAP/Ki67 were performed by sequential incubation of primary antibodies. Fluorochrome conjugated-secondary antibodies, Alexa fluor 568 anti-mouse IgM, Alexa fluor 568 anti-rabbit IgG and Alexa fluor 488 anti-mouse IgG (Molecular Probes, Eugene, Or., USA) were used at 2 µg/ml and incubated for 1 h at room temperature with Hoechst. For monolayer cultures, all the images were obtained using Zeiss AXIO-Observer Z1 microscope (Carl Zeiss SAS, Marly-le-Roi, France). For sphere experiments, images were obtained using Zeiss Lsm 800 airyscan confocal microscope. Images were processed with ImageJ software.

### Self-renewal analysis

After 24 h, cells cultivated in monolayer were treated with 1 nM of GDC-0152 or DMSO for 8 days. At the end of treatment, the supernatant was removed, cells were harvested, dissociated into single cells and plated in 96-well plates (1–5 cells/well) with serum-free medium supplemented with EGF, bFGF and B27. Eight days later, the number of spheres was counted and divided by the original number of cells seeded to evaluate the sphere formation property.

### Cell viability assay

Cells were seeded on poly-DL-ornithin-coated 96-well plates (1500 cells/well). After 24 h, cells were treated with serial concentrations of GDC-0152 (0.01 nM; 1 nM; 100 nM; 1 µM; 10 µM) in 100 μL of cell-specific medium per well for 8 days. After treatment, 10 μl of MTT reagent (3-(4,5-dimethylthiazol-2yl)-diphenyl tetrazolium bromide, Sigma-Aldrich) were added to each well and plates were incubated for 4 h at 37°C. The reduced formazan was dissolved in 100 µL of DMSO and absorbance was measured at 562 nm with an Elx800 microplate reader (Bio-Tek, Colmar, France) and data were analyzed with the Gen5 1.09 software.

### TOF-SIMS imaging

The TOF-SIMS analyses were performed on a TOF V spectrometer (ION-TOF GmbH, Munster Germany) located at Tescan Analytic (Tescan Analytic SAS, Fuveau, France). This spectrometer is equipped with a bismuth liquid metal ion gun (LMIG): 25 keV Bi3+ clusters ions were used for all experiments and an angle of incidence of 45° with respect to the sample surface. The secondary ions were extracted at 2 keV in a single stage reflector time of flight mass spectrometer. Secondary ions were post accelerated to 10 keV at the entrance surface of the hybrid detector, made of one single microchannel plate, followed by a scintillator and a photomultiplier. The data were acquired and processed with SurfaceLab 6.5 software (ION-TOF, GmbH, Munster, Germany). Internal mass calibration was realized using H^+^, H2^+^ and CH3^+^ ions in the positive ion mode and with H^−^, C^−^, CH^−^, CH2^−^ ions in the negative ion mode. Spheres were analyzed following the O^−^ ions. The intensity line scan of O^−^ ion (m/z = 16), scanning the SIMS image of spheroid, was performed in triplicate.

### MALDI-TOF imaging

Spheres were embedded in Gelatin (Sigma Aldrich) at 17.5 mg/mL and stored at −20°C until use. Frozen samples were sectioned at a thickness of 12 μm and then thaw-mounted on a ITO slide (Bruker Daltonics, Mass., USA). The sections were dried for at least 30 min in a dessicator. Optical images were scanned using Opticlab H850 (Plustek, Taïwan) with three teaching marks for a perfect co-localization of optical images and MSI experiments. Sections were then covered with a 2.5-DHB matrix (Bruker Daltonics) and sprayed with TM-Sprayer (HTX Imaging, C., USA). MALDI images were obtained using MALDI-TOF UltrafleXtreme (Bruker Daltonics). FlexImaging 3.4 software (Bruker Daltonics) was used for acquisition and SCILSlab 2016 (SCILS, Bremen, Germany) was used for visualization. The images were normalized using the Root Mean Square normalization. The accurate m/z was chosen thanks to the m/z obtained from 5 spots at different concentrations from 500 nM to 5 µM of GDC-0152 also spotted on the ITO slide.

### PKH67 staining

GBM9 cells were fluorescently marked using a lipophilic dye PKH67 green Fluorescent Cell Linker Kit (Sigma-Aldrich). This staining ensures the monitoring of cell proliferation in cultures for a long period. After 2 washes in PBS, the cell pellet was resuspended in 1 mL of Diluent C and 1 mL of Dye Solution for 2 min. Cells were washed twice with 10 mL of medium, counted and put back in culture.

### Caspase-3 cleavage assay

Spheres were incubated with the green fluorescent NucView™ 488 caspase-3 substrate for profiling caspase-3 activity in living cells (Apoptosis Assay Kit NucView™ 488, Biotium, Inc., Calif., USA), following the manufacturer’s instructions.

### Kinome assay

Cells were treated for 2 h with 1 nM of GDC-0152 or DMSO and extracted in M-PER buffer (Thermo-Scientific) containing inhibitor cocktails. Ten micrograms of total amount of protein with 100 μM of ATP were loaded with fluorescein isothiocyanate (FITC) labeled anti-phospho Serine/Threonine antibodies (Pamgene International B.V.,’s Hertogenbosch, The Netherlands) on PamChip®. Phosphorylation activity was tagged by a FITC-conjugated antibody and recorded with a Pamstation®12 (Pamgene). Results were analyzed with BioNavigator (Pamgene). The experiment was performed in triplicate.

### Statistical analyses

mRNA expression values from microarrays and Q-RT-PCR results were analyzed by the non-parametric Mann-Whitney test. The non-parametric Wilcoxon test was used to analyze several parameters by comparing the effect of GDC-0152 treatment on GB stem-like cell lines in normoxic *versus* hypoxic conditions compared to DMSO-treated control cell lines. All statistical tests were two-sided and the threshold for statistical significance was *P*<0.05. Tests were conducted using the XLSTAT 2013 software (Addinsoft, Paris, France).

## Acknowledgments

This study was supported by Institut National du Cancer (Grants INCa-DGOS-Inserm 6038 and PLBIO N°2014-165), Cancéropôle PACA and institutional grants (Inserm, Aix-Marseille University). A Soubéran is supported by the Association pour la Recherche sur les Tumeurs Cérébrales (ARTC-Sud). We thank P. Weber for confocal microscopy support, L. Dubrez-Daloz and. C. Delfino for technical advices, and R. Pruss for proof reading and suggestions.

## Author’s contribution

AT, AS and DFB conceived the study and wrote the manuscript with the help of GR. VQ developed RNS175 GB stem-like cell line. AS, AT and JC performed most experiments with the help of NBK. CC performed analysis from NCBI GEO data base. JR performed TOF-SIMS analyses. MC and HK performed and analyzed kinomic analyses. CN and DL performed MALDI experiments. All authors discussed the results and commented on the manuscript.

## Synopsis

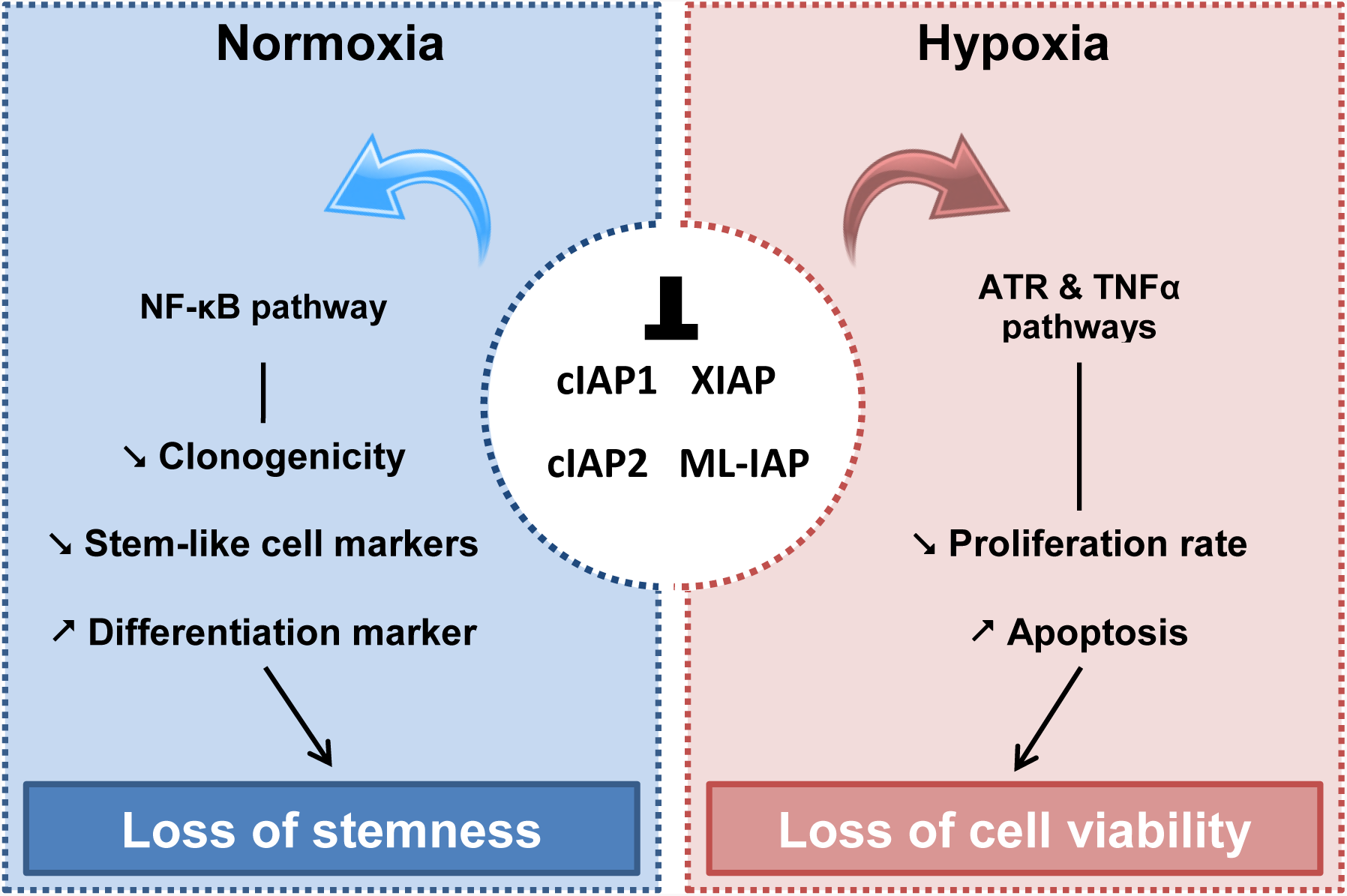

Inhibitor of apoptosis proteins are often overexpressed in cancers including glioblastomas and contribute to treatment resistance. Their pharmacological inhibition using Smac mimetic GDC-0152 determines glioblastoma stem-like cells fate in an oxygen dependent-manner.

- In normoxia, inhibitor of apoptosis proteins inhibition impairs stemness
- In hypoxia, this inhibition induces a loss of cell viability by decreasing cell proliferation and increasing apoptosis.
- These differential effects are subtended by different signaling pathways: NF-κB in normoxia, ATR and TNFα in hypoxia.

**For more informations:** http://www.artcsud.fr/

## Expand view

### Figures legend

**Figure EV1. IAPs distribution between peri-necrotic and non necrotic areas in human glioblastomas and set up of *in vitro* hypoxic condition.**

(**A**) Scatter plots represent cIAP1, cIAP2, XIAP and ML-IAP mRNA expression in palisading cells and common tumor cells of human glioblastomas. Median expression is represented by the horizontal red line and mean expression by the red cross. Data were obtained from NCBI GEO. (**B**) Adrenomedullin (ADM) mRNA was quantified in palisading cells and in common tumor cells. Median expression is represented by the horizontal red line and mean expression by the red cross. Data were obtained from NCBI GEO. (**C**) GBM9 cells were grown in monolayer in normoxia or in hypoxia for 8 days. ADM mRNA level was analyzed by Q-RT-PCR and fold increase ADM mRNA level is shown + SEM (*n*=3 independent experiments). \**P* < 0.05. (**D**) cIAP1, cIAP2, XIAP and ML-IAP mRNA levels of GBM9 cells (grown in monolayer) were analyzed by Q-RT-PCR in normoxia or in hypoxia and fold increase mRNA levels is shown + SEM (*n*=3 independent experiments). ns: not significant.

**Figure EV2. GDC-0152 effect on IAP proteins expression.**

Protein expression levels of cIAP1, cIAP2, XIAP and ML-IAP were analyzed by western blotting. GBM6, GBM9, RNS175 and GBM40 cells were grown in monolayer and were treated either with vehicle alone (DMSO) or with 0.01 nM and 1 nM of GDC-0152 for 8 days in normoxia or in hypoxia. Expression level of β-actin served as loading control.

Normoxia: 20% O_2_ and hypoxia: 2% O_2_.

**Figure EV3. IAPs inhibition triggers loss of stemness in normoxia and decreases cell proliferation and increases cell death in hypoxia.**

(**A-D, F**) GBM6, RNS175 and GBM40 cells were grown in monolayer and treated with vehicle alone (DMSO) or 1 nM of GDC-0152 in normoxia or in hypoxia for 8 days. (**A**) The percentage of self-renewal was calculated as the number of spheres formed divided by the number of cells seeded. Mean + SEM (n=3 independent experiments) is shown. (**B-D**) After treatment, cells were dissociated and stained either with A2B5, anti-CD133 or anti-GFAP (n=6, GBM6; n=8, RNS175; n=4, GBM40) antibodies for flow cytometry analyses. Mean + SEM is shown. (**E**) GBM6, RNS175 and GBM40 cells were treated in monolayer with increasing concentrations of GDC-0152 [0.01 nM; 1 nM; 100 nM; 1 µM; 10 µM] for 8 days in normoxia (red) or hypoxia (blue). Cell viability was expressed as fold increase of controls + SEM (n=4 in triplicate). (**F**) After treatment, cells were dissociated, fixed and stained with Ki67-antibody for flow cytometry analyses. Data are expressed as mean + SEM (n=4). (**G**) DNA fragmentation (SubG0/G1) of DMSO and GDC-0152-treated cells were determined by flow cytometry and percentage of apoptosis is shown. Data are expressed as mean + SEM (n=5). Normoxia: 20% O2 and hypoxia: 2% O2. \**P*<0.05; \*\**P*<0.01; \*\*\**P*<0.001; ns: not significant.

**Figure EV4. Characterization of a glioblastoma 3D model**

(**A**) GBM9 cells were stained with PKH67 and cultivated in suspension in normoxia (20% O_2_). After 8 days, spheres were analyzed by microscopy. Scale bar, 50 µm. A representative staining of 3 experiments is shown. (**B**) GBM9 cells were grown in suspension for 8 days in normoxia (20% O_2_). After 8 days GFAP and Ki67 immunofluorescences were performed and counterstained with Hoechst. Localization of GFAP^+^ cells (red) and Ki67^+^ cells (green) in GBM9 spheres was analyzed by confocal microscopy on 15 z stacks projection. For the top of the sphere (up) 1 to 5 z were projected, for middle up 6 to 10 z and for middle down 11 to 15 z were projected. Representative stainings of 3 independent experiments are shown. Scale bar, 50 µm. (**C**) Eight-days old spheres were treated with vehicle alone (DMSO) or 1 µM of GDC-0152 for 8 days. Spheres (n=10-12), were localized using the phosphocholine signal (m/z 184.8) using MALDI-TOF imaging, and images of GDC-0152 (m/z 499.4) and sodium adduct (m/z 521.4) were analyzed. Spheres appeared in black on the scan images, colored in red by the phosphocholine signal and the GDC-0152 in green. Scale bar, 1.1 mm.

**Figure EV5. Analysis of Serine threonine kinases activation upon IAPs inhibition depending on oxygen level**

(**A-B**) GBM9 stem-like cells were cultivated in monolayer in normoxia (**A**) or hypoxia (**B**) and treated with vehicle alone (DMSO) or 1 nM of GDC-0152 for 2 h. Serine/threonine kinases activity was analyzed by PamGen kinome assay. On graphs are listed the main kinases either more (negative values) or less (positive values) activated in GDC-0152- treated cells compared to DMSO-treated cells. (**C**) Table summarizes the main kinases activated with GDC-0152 treatment compared to vehicle, in normoxia and in hypoxia.

